# SeverYX: An ensemble framework for robust variant severity classification

**DOI:** 10.64898/2026.11.03.762814

**Authors:** Evyatar Mor, Eldar Dedic, Yaron Mazor, Suzanne Drury, Hagar Mor-Shaked, Lina Zelinger

**Affiliations:** Geneyx Genomex LTD, Herzliya, Israel

**Keywords:** Variant Prediction, Clinical Genomics, Ensemble approach, Variant Interpretation, Decision Tree, Computational pathogenicity prediction

## Abstract

Accurate interpretation of genetic variants in clinical genomics depends on reconciling multiple computational predictors that often disagree or provide incomplete evidence. We present SeverYX, a transparent, rule-based ensemble framework that consolidates conservation, splicing, and functional-impact predictors into a three-tier severity classification (‘High’, ‘Medium’, ‘Low’) through an effect-aware decision architecture. We benchmarked SeverYX against a curated ClinVar dataset of 270,208 MANE-transcript variants (86,161 pathogenic/likely pathogenic, 184,047 benign/likely benign), stratified by molecular consequence, review status, and haploinsufficiency. The baseline configuration achieved 97.6% concordance with ClinVar; at a fixed High cut, 90.0% of pathogenic and 2.0% of benign variants were assigned High. Concordance was highest for loss-of-function variants and lowest for non-coding and unclassified pathogenic variants, reflecting known gaps in current predictive tools. Systematic threshold optimization, combined with the addition of AlphaMissense and REVEL as missense predictors, raised overall concordance to 98.7%, reduced the benign High rate to 0.4% (at a cost of pathogenic High sensitivity falling to 88.3%), and improved the Matthews correlation coefficient from 0.895 to 0.908. These results show that SeverYX offers a transparent, reproducible and extensible approach for consolidating heterogeneous variant-effect evidence into clinically interpretable severity calls, and a scalable path for incorporating future predictive models into genomic interpretation workflows.

## Introduction

The continuously increasing use of high throughput sequencing technologies has substantially advanced molecular genetic testing, greatly enhancing both research capabilities and the clinical and molecular diagnosis of hereditary conditions [1–3]. To ensure consistent reporting across the genomic analysis ecosystem, American College of Medical Genetics and Genomics (ACMG) and the Association for Molecular Pathology (AMP) have built a framework for the clinical reporting of sequence variants associated with human disease [4], enabling more accurate classification of variant pathogenicity. Despite this, significant challenges remain when translating high throughput sequencing outputs into clinically actionable variant interpretations.

Variant annotation is essential for establishing meaningful and actionable links between the identified genomic variants and reported phenotypic traits, forming the fundamental building block of any interpretive workflows. The accuracy of this process relies on precise variant localization, detailed characterization of molecular consequences, and rigorous assessment of potential impacts on gene and protein function. This process encompasses clinical categorization (e.g. pathogenic vs benign, as seen in ClinVar; https://www.ncbi.nlm.nih.gov/clinvar/), integration of sequence constraint metrics (e.g. GERP [5] and PhyloP [6]), and evaluation of predicted effects on protein structure and molecular behavior (e.g. CADD [7], SIFT [8], REVEL [9], SpliceAI [10], MutationTaster [11], and AlphaMissense [12]). Together, these resources provide complementary evolutionary and functional evidence layers that inform assessments of variant deleteriousness. Yet, because individual algorithms often produce discrepant or partially overlapping predictions [13,14], their interpretation demands expert adjudication, contextual biological insight, and substantial analytical effort. Consequently, a unified framework capable of synthesizing these heterogeneous evidence sources into a coherent, consistent, and interpretable variant impact severity assessment would not only streamline clinical genomics workflows but also reduce inter-analyst variability and enhance the reliability and reproducibility of variant interpretation.

To address this need, we developed SeverYX, a rule-based ensemble work frame designed to assign each variant to one of three severity categories (‘High’, ‘Medium’, ‘Low’). The algorithm organizes the predictive space using explicit thresholds logic, enabling fully reproducible evaluation, modification, and automated optimization.

## Materials and Methods

### ClinVar-based Variant Database Construction and Preprocessing

A ClinVar-based variant database was generated as a benchmark for evaluating SeverYX assignments. The ClinVar GRCh38 VCF release dated 07 Jan 2024 (https://ftp.ncbi.nlm.nih.gov/pub/clinvar/vcf_GRCh38/archive_2.0/2024/clinvar_20240107.vcf.gz) was downloaded from the NCBI repository, decompressed, and processed with a dedicated Python pipeline that parses INFO annotations, restricts variants to genes in one of three haploinsufficiency categories (score 0, score 3, or score 40) to capture distinct dosage-sensitivity contexts, and standardizes transcript context by retaining only variants annotated on MANE transcripts to avoid transcript-mapping ambiguity. Within each haploinsufficiency group, variants were assigned with effect prediction classes aligned with SeverYX framework using ClinVar annotations and Variant Effect Predictor (VEP) consequences (Loss-of-Function, missense, splice-region coding, start/stop, and other), then stratified by ClinVar clinical significance (pathogenic/likely pathogenic vs benign/likely benign) and by simplified review status (one-star vs two-star or more). Variants annotated as uncertain significance or exhibiting incomplete/inconsistent annotation patterns were excluded. After filtering, 270,208 variants remained across all haploinsufficiency groups and effect classes; the final dataset was exported to a standard VCF (Supplementary Fig. 1).

### SeverYX algorithm construction and optimization strategy

SeverYX assigns to each variant one of three severity labels (‘High’, ‘Medium’, or ‘Low’) using a rule-based decision framework that integrates 11 predictive scores (Table 1, 9 initially and two additional predictors were added later as part of the optimization and framework construction) and variant effect annotations. The SeverYX optimization algorithm systematically enumerates candidate threshold configurations by sweeping each field’s value range (defined by min, max, and step size) and its allowed logical operators. For every candidate configuration, a deep copy of the base decision-tree logic is serialized as an isolated JSON file, ensuring thread-safe parallel evaluation across available CPU cores. Each configuration is independently applied to the full variant dataset, producing evaluation metrics - the observed success ratio and balanced accuracy, defined as (sensitivity + specificity)/2-which are used to identify the optimally performing threshold set. The JSON-based decision-tree structure of the configurations ensures full reproducibility and auditability of every combination of parameters. The code and implementation plan are available at https://github.com/geneyx/geneyx.severity.

**Table 1.**
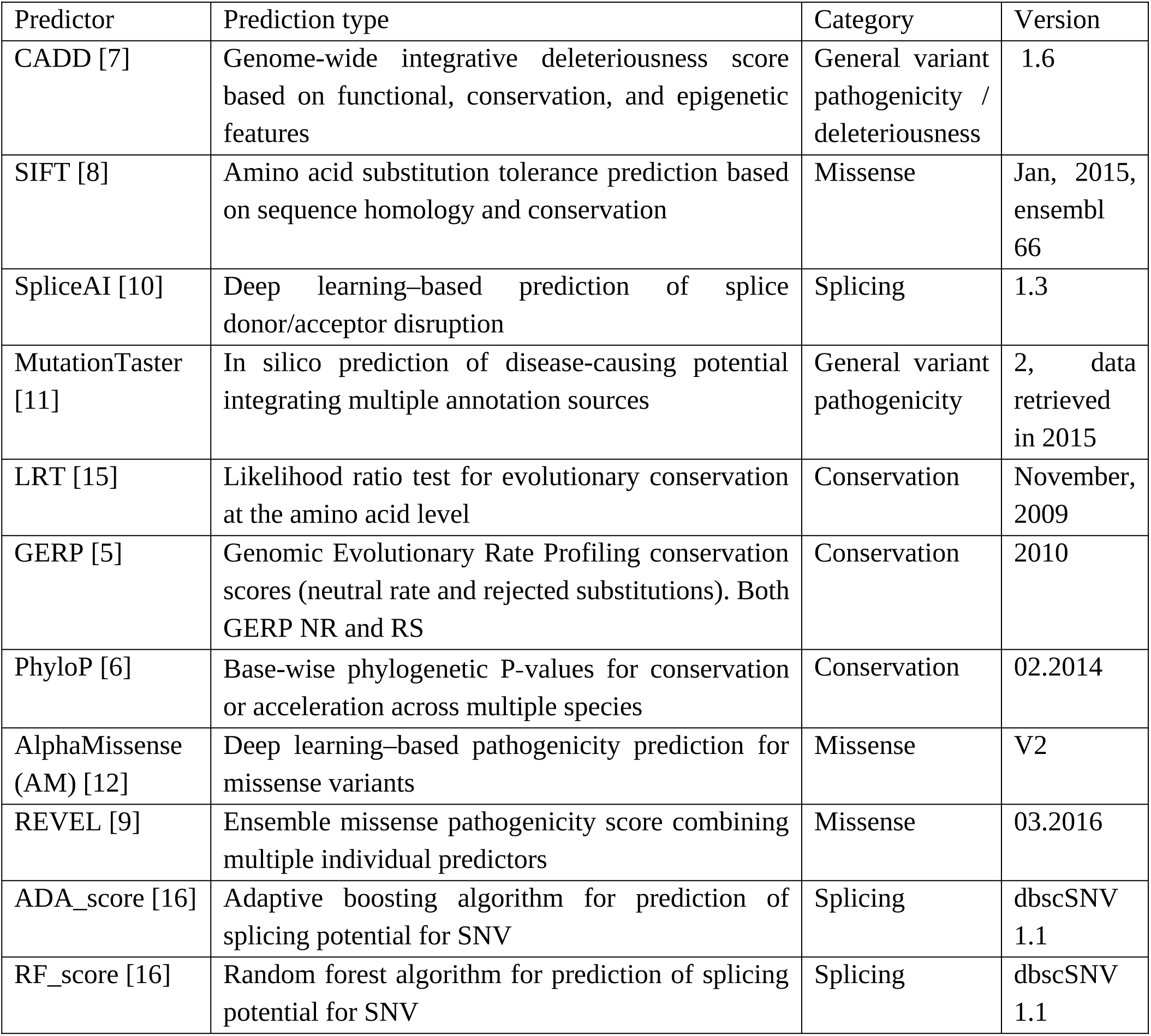
Predictors used by SeverYX for severity category determination.

Each rule in the decision tree specifies a predictor, a comparison operator, and a threshold value that was determined initially based on established guidelines and published literature for each predictive tool. Rules are evaluated in a fixed, short-circuited hierarchy, yielding an effect-aware severity assignment.

Effect classes for variants are derived from standard and established molecular consequence terms. Nonsense, frameshift and canonical splice-site (donor and acceptor) variants constitute the loss of function (LoF) group; non-synonymous coding variants define the Missense group; start-gain, start-loss, and stop-loss variants define the Start/Stop group; Splice-region coding Effect is defined by non-synonymous coding and splice site region or synonymous coding and splice site region variant and remaining variants are assigned to the Other group.

Evaluation proceeds in several stages. First, global high-severity rules test for combinations of strongly abnormal scores across two predictors. Variants satisfying the standalone high-severity requirement (Table 2) are classified as ’High‘ severity and are not processed further. Then, variants that remain unclassified are evaluated within their functional effect class; LoF, Missense, Splice-region coding, Start/Stop, or Other, using effect specific high and medium rules that combine one or more predictors with effect-appropriate thresholds (Table 3 and Supplementary Table 1). Subsequently, standalone rules for ‘Medium’ classification are applied. Finally, variants not meeting any ‘High’ or ’Medium‘ criteria are assigned ’Low‘ severity, indicating limited or discordant evidence for deleterious impact.

**Table 2.** Standalone Thresholds.

| **Predictor** | **Severity Assignment** | **Baseline threshold<sup>#</sup>** | **Threshold post optimization<sup>\$</sup>** |
|---|---|---|---|
| AdaScore | Medium | > 0.6 | > 0.6 |
|  | High | > 0.6 | NA |
| CADD (PHRED) | High | ≥ 25.0 | ≥ 27.0 |
|  | Medium | ≥ 25.0 | ≥ 25.0 |
| RfScore | Medium | > 0.6 | > 0.6 |
|  | High | > 0.6 | NA |
| SpliceAI | High | ≥ 0.9 | ≥ 0.85 |
|  | Medium | ≥ 0.9 | ≥ 0.9 |

\# In the baseline configuration the ‘High’ severity requirement was any two of four criteria. <sup>$</sup>In the optimized configuration ‘High’-severity requirement is satisfied only when both the SpliceAI and the CADD condition are met; and ‘Medium’ severity requirement is satisfied by any one of the four conditions listed.

**Table 3.**
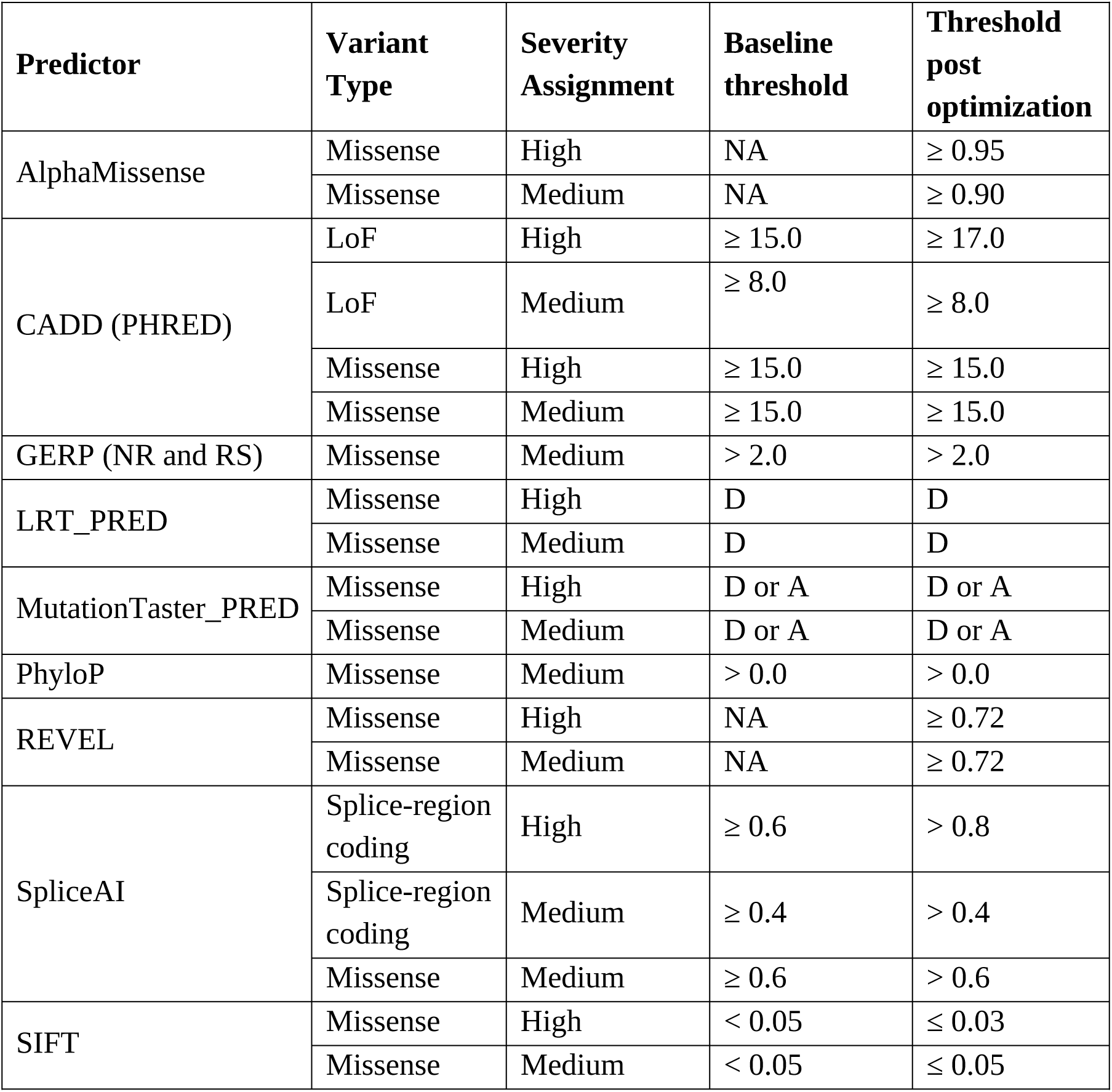
Combination Thresholds.

### Analysis and validation

The processed reference VCF was uploaded to the Geneyx platform. For each variant, the pipeline produced the severity classification, the predictor scores used in the severity decision tree, and ClinVar-derived annotation fields, including clinical significance and review status. The resulting annotated table was exported in tab-delimited format and served as input to a custom validation program.

The validation program implements a structured evaluation pipeline. First, variants are grouped according to haploinsufficiency category, functional effect class, clinical significance, and review status, using the same definitions applied during database construction. Variants lacking a Geneyx severity label or missing essential ClinVar attributes for subgrouping are excluded. The program then classifies each variant as “expected” or “unexpected” based on predefined assumptions: pathogenic and likely pathogenic variants are expected to receive ‘Medium’ or ‘High’ severity assignment, and benign and likely benign variants are expected to receive ’Low‘ or ’Medium‘ severity assignment. Pathogenic or likely pathogenic variants with ’Low‘ severity and benign or likely benign variants with ’High‘ severity are designated unexpected; all other combinations are designated expected. These were used to calculate the base concordance rate between SeverYX and ClinVar designation.

From these expected/unexpected classifications, formal performance measures are derived: pathogenic and likely pathogenic variants with an expected assignment (‘High’ or ‘Medium’) are treated as true positives and benign and likely benign variants with an expected assignment (‘Low’ or ‘Medium’) as true negatives, while benign and likely benign variants assigned ‘High’ are false positives and pathogenic and likely pathogenic variants assigned ‘Low’ are false negatives. True positive and false-positive rates are computed from these quantities, alongside the global success ratios reported above. Because ‘Medium’ is counted as an expected assignment for both classes, these two rates are not measured at a common decision threshold: the true positive rate is evaluated at a ‘Medium or above’ cut, whereas the false-positive rate is evaluated at a ‘High’ cut, so the pair does not define a single achievable operating point. Operating characteristics are therefore also reported at each of the two internally consistent cuts, ‘High’ against ‘Medium or Low’ and ‘High or Medium’ against ‘Low’, and the complete cross-tabulation of severity against clinical significance is given for both configurations in the Results.

The same framework is used to assess and refine SeverYX thresholds. The validation program reads the configuration file, recomputes severity labels under alternative threshold settings, and identifies variants whose classifications change relative to the reference configuration. For each candidate configuration, success ratios and confusion matrix derived metrics are recalculated on the full ClinVar-based dataset.

### Statistical analysis

Analyses were performed in Python 3.9 (pandas, NumPy and SciPy). The unit of analysis was the variant; each variant was assigned to exactly one level of each stratifying factor. The primary endpoint was the concordance rate, the proportion of variants with an expected severity assignment, estimated overall and within each haploinsufficiency, effect-class, clinical-significance and review-status stratum, with two-sided 95% Wilson score confidence intervals. Secondary endpoints, derived from the 2 × 2 table cross-classifying clinical significance against expected/unexpected assignment, were sensitivity, specificity, false-positive rate, balanced accuracy ((sensitivity + specificity)/2), the Matthews correlation coefficient and Cohen’s κ. Because SeverYX returns an ordinal label rather than a continuous score, no continuous-score receiver-operating-characteristic curve was estimated; operating characteristics are instead reported at each of the two achievable ordinal cuts, together with the trapezoidal area under those two points. Strata were compared by Pearson’s χZ test (Fisher’s exact test where any expected count was below five) with odds ratios, and baseline versus optimized configurations by McNemar’s exact test. Two-sided p < 0.05 was considered significant.

### Ethics statement

This study did not involve new human subjects research, primary patient contact, or the collection of original clinical samples. All variant data were obtained from ClinVar, a public, de-identified repository of previously submitted variant classifications; no individual-level identifiable data were accessed, and no new biological samples were collected. Accordingly, no institutional review board (IRB) or research ethics committee (REC) review was sought, and no informed consent procedures were applicable. This study did not involve animal subjects.

## Results

### Development of the ensemble algorithm

The SeverYX algorithm was designed to unify diverse predictive and annotation layers into a single, interpretable three-tier severity system (‘High’, ‘Medium’, ‘Low’) aimed at simplifying variant triage without sacrificing analytical depth.

The input to the algorithm is a list of annotated variants produced by Geneyx Analysis. We employed a short-circuited rule-based structure, meaning that evaluation stops at the first satisfied condition, thus favoring computational efficiency and interpretability (Figure 1). Its logic integrates (1) stand-alone High-severity conditions, (2) Medium-effect consequence flags (including codon deletions/insertions and start/stop disruptions), and (3) effect specific evidence blocks tuned independently for loss-of-function (LoF), missense, splice-region coding effect, start/stop loss, and other molecular consequence categories (see Methods section for details). All rules are fully parameterized within a log threshold configuration file (JSON format, can be found in the repository listed under Data & Code Availability) for each prediction tool (Table 1), enabling reproducibility, direct operator specification, and explicit control over null-value handling. This architecture yields a transparent and configurable model in which global and effect-specific rules, anchored in widely used predictors, converge on a reproducible three-tiered severity label suitable for large-scale analysis.

**Figure 1.**
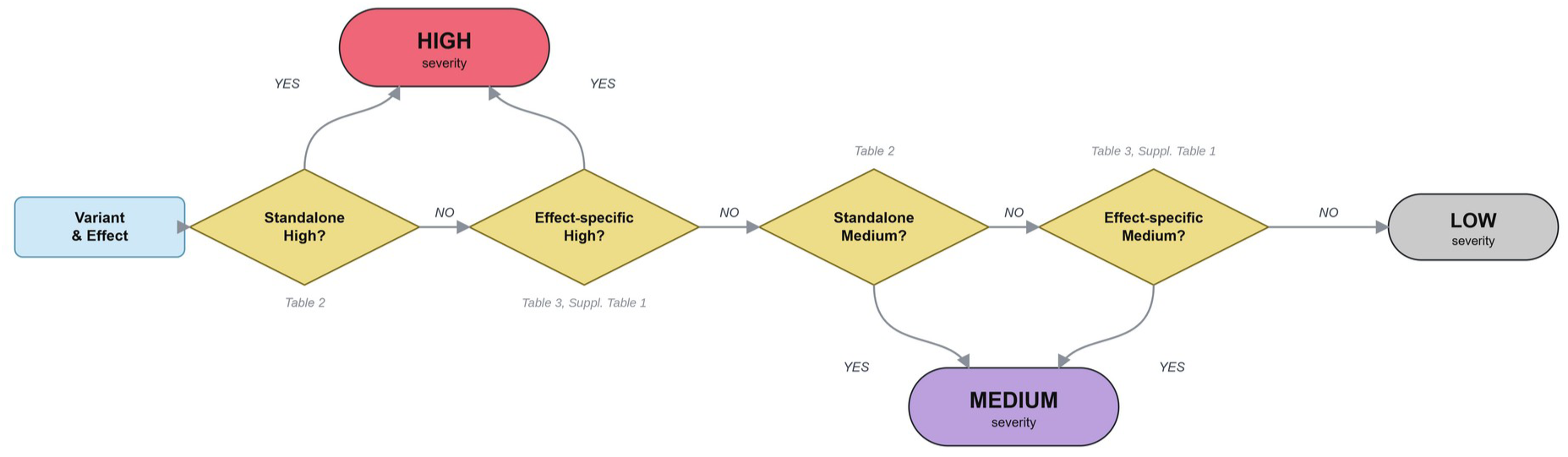
SeverYX dissection workflow. The diagram illustrates a stepwise process for classifying variant severity (‘High’, ‘Medium’, or ‘Low’) using a structured ensemble rule-based framework.

### Validation of Severity Classification Performance Using ClinVar

In order to assess the performance of SeverYX and to benchmark it against public dataset we utilized the ClinVar repository. Our hypothesis was that variants classified as pathogenic/likely pathogenic (P/LP) in ClinVar should not be assigned ’Low’ in the algorithm, and variants classified as benign/likely benign (B/LB) should not be assigned ’High’ severity.

We evaluated 270,208 ClinVar variants (86,161 P/LP, 184,047 B/LB, from three haploinsufficiency categories, Supplementary Figure 3A) using the SeverYX algorithm, each variant receiving one of three possible assignments (Figure 2A), with majority classified as ‘Low’ severity concordant with majority of variants being labeled as “Benign” in ClinVar. Overall concordance between ClinVar and SeverYX classification was 97.6%, CI(97.55– 97.66%), demonstrating that the model captures the majority of ClinVar supported pathogenicity trends (Figure 2B), and concordance varied by no more than 3.0 percentage points across haploinsufficiency groups (98.51%, CI (98.39-98.62), 97.44%, CI (97.37-97.50) and 95.47%, CI (94.05-96.57) for the no-evidence, sufficient-evidence and dosage-sensitivity-unlikely groups respectively; χZ(2) = 202.0, p < 0.001; Supplementary Figure 3B), although the dosage-sensitivity-unlikely group (n = 1,060) was estimated with correspondingly wider uncertainty.

**Figure 2.**
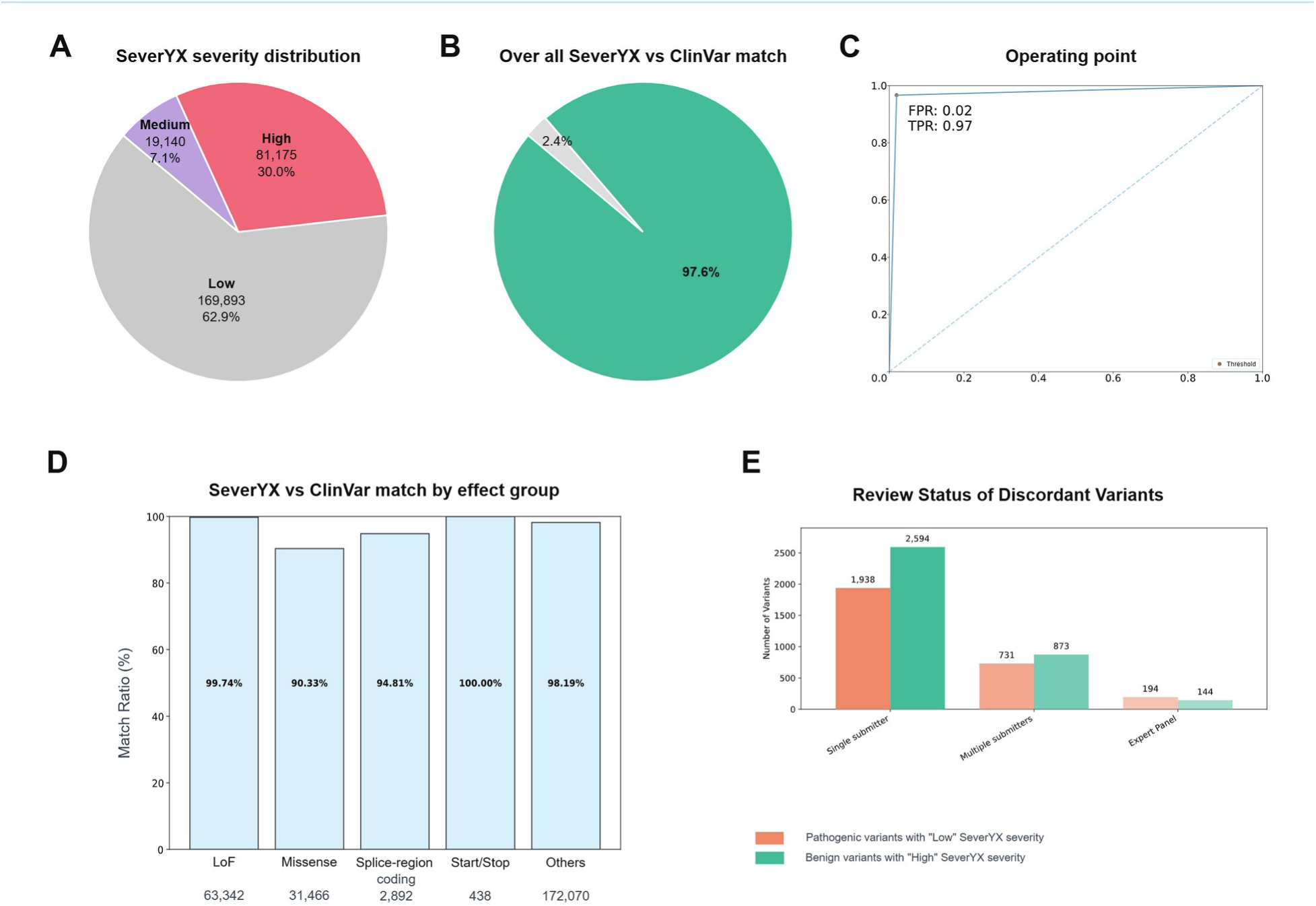
Performance assessment of the SeverYX severity-scoring framework using ClinVar dataset. (A) Distribution of SeverYX severity scores across all evaluated variants, categorized into ‘High’, ‘Medium’, and ‘Low’ severity groups. (B) Overall concordance between SeverYX severity assignments and ClinVar pathogenicity categories, showing the proportion of matched versus mismatched annotations. (C) SeverYX operating point in receiver-operating-characteristic space relative to ClinVar. Because SeverYX returns an ordinal severity assignment rather than a continuous score, a single (false-positive rate, sensitivity) point is defined; the connecting segments are interpolations to (0,0) and (1,1) and do not represent a fitted ROC curve. The false-positive rate shown (0.02) is measured at a ‘High’ cut and the sensitivity shown (0.97) at a ‘High or Medium’ cut, so the two coordinates are not measured at the same decision threshold; operating characteristics at each internally consistent cut are given in the Results. (D) Match rates between SeverYX and ClinVar stratified by variant effect groups, including loss-of-function, missense, splice-region coding, start/stop variants, and other categories. Number of variants in each group are listed below each bar. (E) Number of discordant variants by ClinVar submitter category (single submitter, multiple submitters, expert panel), shown separately for P/LP variants assigned ‘Low’ (1,938, 731 and 194) and B/LB variants assigned ‘High’ (2,594, 873 and 144). Counts are not normalised to subgroup size: discordance rates were 2.39% for one-star and 2.40% for two-star-or-better entries.

Cross-tabulating severity against clinical significance gave, for P/LP variants, 77,564 ‘High’, 5,734 ‘Medium’ and 2,863 ‘Low’, and for B/LB variants 3,611 ‘High’, 13,406 ‘Medium’ and 167,030 ‘Low’. At a fixed ‘High’ cut this corresponds to a sensitivity of 0.900 and a false-positive rate of 0.020 (balanced accuracy 0.940); at a ‘High or Medium’ cut, to a sensitivity of 0.967 and a false-positive rate of 0.092 (balanced accuracy 0.937). The two coordinates plotted in Figure 2C are measured at these two different cuts and therefore do not describe a single achievable operating point. At the fixed ‘High’ cut the Matthews correlation coefficient was 0.895 and Cohen’s κ 0.894; the trapezoidal area under the two achievable operating points was 0.969. The model assigned an expected severity to 83,298 P/LP variants (true positives) and to 180,436 B/LB variants (true negatives). Because the 19,140 variants (7.1%) assigned ‘Medium’ (Figure 2A) are counted as expected for both classes, we also report a strict criterion counting only ‘High’ as concordant for P/LP and only ‘Low’ for B/LB, under which concordance was 90.5%. Conversely, 3,611 benign variants received ’High‘ severity (false positives), and 2,863 pathogenic variants received ’Low‘ severity (false negatives).

Analysis across effect groups revealed consistently high predictive alignment: the LoF group achieved the highest concordance among the classes with effect specific rules, at 99.74%, followed by the other (98.19%), splice-region coding-effect (94.81%) and missense (90.33%) groups (Figure 2D). All Start/Stop variants are mapped to ‘Medium’ severity by design, so the 100% concordance of that group reflects this construction rather than predictive performance. The LoF aggregate is similarly constrained: because the LoF high-severity rule is satisfied when CADD is missing, almost all LoF variants receive ‘High’, so the 99.74% figure largely reflects the fact that 99.7% of that class is P/LP rather than discrimination within it. Concordance also differed sharply by clinical significance within the LoF group: all 63,177 P/LP LoF variants received an expected assignment, whereas none of the 165 B/LB LoF variants did, because the LoF high-severity rule deliberately treats a missing CADD score as satisfying its threshold. In contrast, the ‘Other’ P/LP subgroup reached only 56.03% concordance, reflecting reduced predictive resolution for the heterogeneous non-coding, synonymous and unclassified consequences aggregated in this class. Conversely the B/LB ‘Other’ subgroup, 165,588 variants or 61.3% of the benchmark, is 100.00% concordant because variants of these consequences match no effect specific rule and are assigned ‘Low’ by default; benign ‘Low’ assignments alone account for 63.3% of all concordant calls, so the overall concordance figure is dominated by this group. The full subgroup-level breakdown, combining effect class, clinical significance, review status and haploinsufficiency group, is given in Supplementary Figure 2.

Discordant classifications were most numerous among variants with single-submitter review status (Figure 2E), in proportion to their share of the benchmark; discordance rates were similar across review-status strata (one star, 4,534/189,328 = 2.39%; two stars or more, 1,940/80,880 = 2.40%; χZ(1) = 0.002, p = 0.96, odds ratio 1.00). Together, these findings demonstrate strong alignment between SeverYX outputs and curated ClinVar data while highlighting known limitations in low-evidence or non-coding domains.

### Optimization of Severity Thresholds to Improve Predictive Performance

To refine classification performance beyond the baseline rule set, we conducted systematic threshold optimization using the configurable JSON severity logic. Thresholds at seven rule positions, involving four predictors that participate in the high-severity rules (SpliceAI, CADD, AM and SIFT), were swept across pre-specified ranges: SpliceAI 0.80–0.95 in steps of 0.05 for the stand-alone rule and 0.4–0.8 in steps of 0.2 for the splice rule, CADD 21–27, 11–17 and 11– 15 in steps of 2, AM 0.80–0.95 in steps of 0.05, and SIFT 0.03–0.07 in steps of 0.02, enumerating 6,912 configurations of which 4,295 were evaluated. Six of the seven selected values lie at the edge of their range, so the grid is truncated at its own optimum and a better configuration may lie outside it. Each resulting configuration was scored on the full reference dataset for overall success ratio, sensitivity, false-positive rate and balanced accuracy, with parallel execution across available CPU cores enabling efficient screening of thousands of candidate configurations.

This search (Supplementary Figure 4) identified the threshold set that scored highest within the searched grid. Several cutoffs moved toward more discriminative values: the loss-of-function CADD requirement rose from ≥15 to ≥17, the splice site SpliceAI threshold rose from ≥0.6 to >0.8, and the missense SIFT threshold tightened from ≤0.05 to ≤0.03 (Table 2, Table 3). The stand-alone high-severity rule was also simplified, from any two of four conditions (SpliceAI, CADD, AdaScore and RfScore) to both of two conditions, with SpliceAI relaxed slightly (≥0.9 to ≥0.85) and CADD raised (≥25 to ≥27). Applied together, these changes moved the operating point leftward in ROC space, the false-positive rate at a fixed ‘High’ cut falling from 0.020 to 0.004 while sensitivity at that same cut fell from 0.900 to 0.883, raising overall concordance from 97.6%, CI (97.55–97.66) to 98.7%, CI (98.62–98.71) (Figure 3C). In absolute terms the change traded 2,877 fewer B/LB variants assigned ‘High’ (3,611 to 734) against 1,475 fewer P/LP variants assigned ‘High’ (77,564 to 76,089); balanced accuracy at a fixed ‘High’ cut was 0.940 before and after, and strict concordance was 90.5% and 90.0% respectively, so the gain is largely a shift in the false-positive/true-positive balance; on measures that use both achievable cuts the improvement in discrimination was modest but consistent, the Matthews correlation coefficient rising from 0.895 to 0.908, Cohen’s κ from 0.894 to 0.905 and the trapezoidal area under the two achievable operating points from 0.969 to 0.976. Absolute values in Supplementary Figure 4 are not directly comparable with those in Figure 3A because the threshold sweep was performed on the rule set that preceded REVEL integration. That sweep configuration is a third rule set, distinct from both the baseline and the optimized configuration reported here: it included AM but not REVEL, and its loss-of-function medium rule required two of a single condition and was therefore unsatisfiable. Supplementary Figure 4 accordingly ranks threshold sets within that historical rule set and is not a comparison with either reported configuration. Adjustments concentrated on loss-of-function and splice-related predictors produced more stable classification regions; the variants reassigned away from ‘High’ moved almost entirely to ‘Medium’ rather than to ‘Low’ (2,863 versus 2,864 P/LP variants assigned ‘Low’), so sensitivity at a ‘High or Medium’ cut was unchanged.

**Figure 3.**
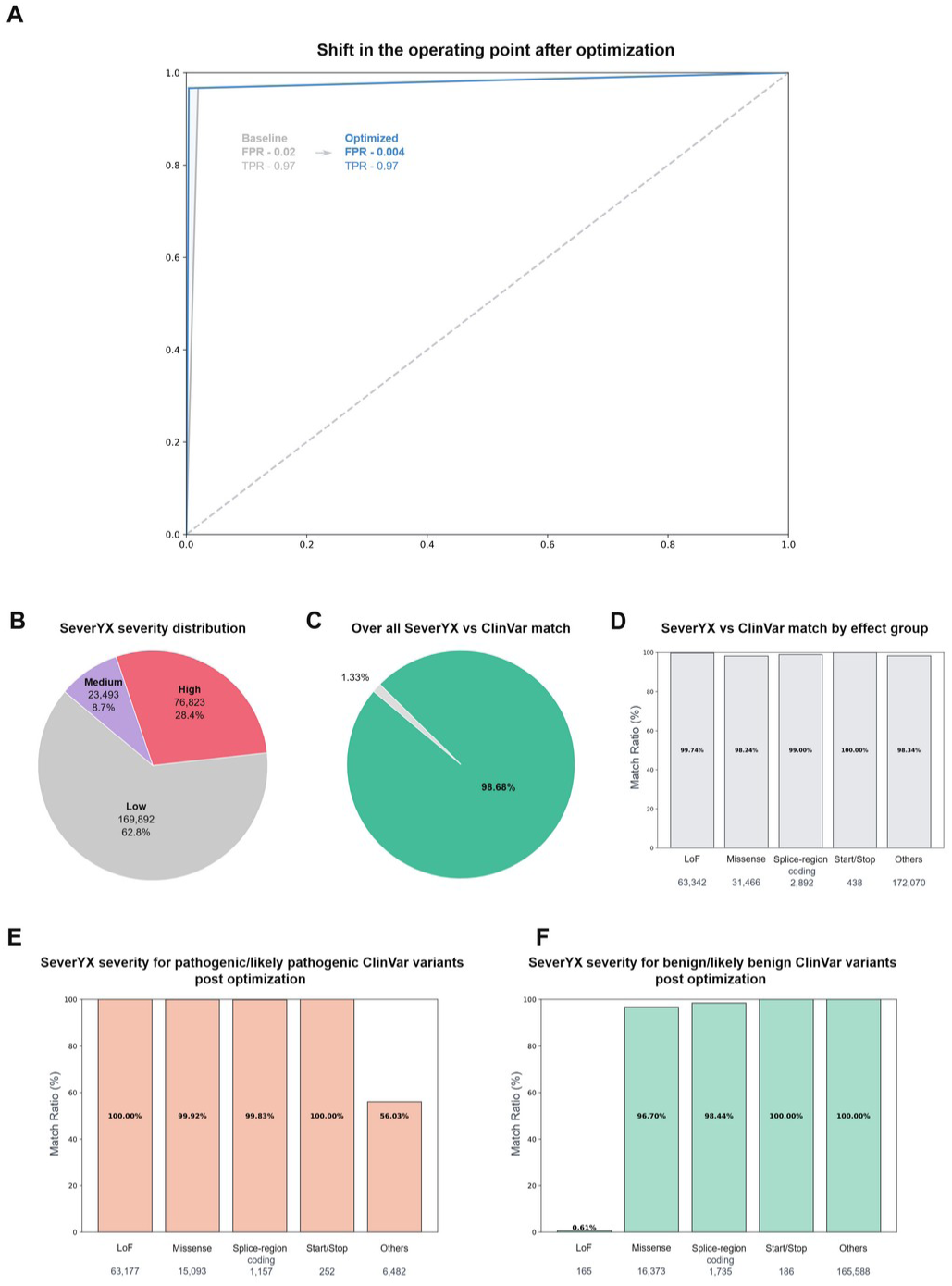
SeverYX optimization and post-optimization performance. (A) Comparison of operating points before and after threshold optimization. The panel shows the shift in false-positive rate (FPR) and in true-positive rate (TPR) between the baseline and optimized configurations. The optimized rule set moves the operating point leftward, that is to a lower false-positive rate at unchanged true-positive rate; the marked point is the selected decision threshold used in downstream analyses. FPR is measured at a ‘High’ cut whereas TPR is measured at a ‘High or Medium’ cut, so the two coordinates are not measured at the same decision threshold (see Methods and Results). FPR, false-positive rate; TPR, true-positive rate (sensitivity). (B) Distribution of optimized SeverYX severity scores across all analysed variants, showing the proportions classified as High, Medium, and Low severity. (C) Overall concordance between SeverYX severity assignments and ClinVar clinical significance labels following optimization, indicating the percentage of matching versus non-matching variants. (D) Match rates between SeverYX and ClinVar stratified by variant effect groups, including loss-of-function, missense, splice-region coding, start/stop variants, and other categories; sample sizes for each group are displayed below the bars. (E) Concordance between optimized SeverYX severity scores and ClinVar designations specifically for variants annotated as *pathogenic or likely pathogenic* in ClinVar, stratified by effect category. (F) Concordance between optimized SeverYX severity scores and ClinVar annotations for variants labeled *benign or likely benign* in ClinVar, also stratified by effect category. Together, these panels illustrate the shift in operating characteristics achieved by threshold optimization and the resulting alignment between SeverYX severity outputs and ClinVar reference labels across multiple variant classes.

### Expanding the Severity Framework with AlphaMissense and REVEL

As predictive modeling advances, incorporating new information and predictor tools becomes essential for accurate variant interpretation. We expanded SeverYX by integrating AM and REVEL, two state-of-the-art predictors for missense pathogenicity. Analysis of AM distributions revealed clear bimodality in expected variants: P/LP missense variants clustered tightly near 1.0, whereas B/LB variants were concentrated near 0.1. Unexpected variants showed that P/LP clustering near 1.0 was preserved, but benign classifications exhibited broader variability, suggesting that AM reliably identifies deleterious missense changes but may be less conservative for benign outcomes. Although the pathogenicity threshold suggested in the original AM publication is 0.564 [12], our initial configurations used AM ≥0.89, optimization repeatedly identified AM ≥0.95 as a superior discriminator between pathogenic and benign missense variants, supported by distinctly bimodal AM score distributions (Supplementary Figure 5A). REVEL distributions similarly aligned with expected ClinVar groupings, reinforcing its value as a complementary predictor. Combining AM and REVEL within the missense decision path substantially improved classification specificity, particularly when both tools independently supported high impact predictions. These enhancements were integrated into the updated severity flow, which now incorporates both stand-alone and effect specific rules influenced by the new predictors.

Performance analytics post integration of the optimized configuration incorporating both predictors (Figure 3B–D) showed that missense concordance rose from 90.33% to 98.24% and overall concordance to 98.7%. Stratified by clinical significance, all P/LP LoF and Start/Stop variants and ≥99.8% of P/LP missense and splice-region coding-effect variants received an expected assignment, while P/LP ‘Other’ variants remained at 56.03% (Figure 3E); among B/LB variants concordance was ≥96.7% in every group except LoF, where 1 of 165 variants received an expected assignment (Figure 3F), for the reason given above. Because threshold optimization and the addition of the two predictors were applied together, their individual contributions cannot be separated from these data. Moreover, the gains were most pronounced in variant subsets where predictor ambiguity previously limited discrimination accuracy, such as borderline missense cases that lacked decisive support from earlier predictor tools such as SIFT or CADD (Supplementary Figure 5).

Overall, extending the severity decision framework with AM and REVEL significantly strengthened the algorithm’s ability to capture functional missense consequences, demonstrating the adaptability of SeverYX to emerging genomic prediction technologies.

## Discussion

Evaluating variant impact from high-throughput sequencing remains a persistent challenge in clinical genomics, particularly for variants lacking confirmatory functional studies. Although numerous computational tools exist to predict variant impact, they vary widely in scope, many assess only specific variant classes, and they frequently produce conflicting or only partially overlapping predictions. This inconsistency is well documented: Smith and Kitzman showed that commonly used splicing predictors disagree substantially when benchmarked against massively parallel splicing assays [14], and Chen et al. reported only 58.5% transcript-level concordance among ANNOVAR, VEP and SnpEff, with substantial disagreement on loss-of-function calls [13]. As a result, analysts are required to repeatedly reconcile, consolidate, and update variant impact annotations, a time-intensive process prone to inter-reviewer variability.

SeverYX addresses this by consolidating multi-feature genetic evidence from tools such as CADD, SIFT, REVEL, MutationTaster, SpliceAI, and conservation metrics (GERP, PhyloP) into a single, explicit three-tier severity framework. By collapsing heterogeneous signals into structured decision gates, the system reduces the conflicts that arise when individual tools disagree. This approach aligns with the growing recognition within the community that computational evidence must be used carefully and transparently: the ClinGen SVI Working Group has emphasized the need for calibration and evidence-strength mapping for computational predictors within ACMG/AMP criteria (PP3/BP4) [17]. SeverYX shares these aims of transparency and explicit thresholds, but departs from the SVI approach by combining several predictors through unweighted condition counts rather than a single calibrated predictor at a likelihood-ratio-derived evidence strength; its severity tiers are therefore not ACMG evidence codes and are not intended to be used directly as PP3 or BP4.

Beyond this design philosophy, SeverYX offers several concrete advantages. Its rules and thresholds are defined entirely in an external, versioned configuration file rather than embedded in model weights, so every decision pathway, not just the final label, is auditable and reproducible. Its effect-specific rule blocks let it operate across heterogeneous variant categories, including coding, splice-related and non-coding contexts, within a single modular architecture, which helps mitigate (though does not eliminate) the reduced predictive resolution documented for non-canonical splice and regulatory variants [14,18,19]. And because its logic is externally configured rather than learned, it is forward-compatible with new prediction tools: as predictors such as AlphaGenome [20] emerge, they can be incorporated and calibrated, following the same approach ClinGen SVI used for AM, ESM1b and VARITY [17], without redesigning the underlying decision architecture.

These design choices translate into strong empirical performance. SeverYX achieved 97.6% baseline concordance with ClinVar, rising to 98.7% after threshold optimization and the addition of AM and REVEL as missense predictors, with the largest gains concentrated in the previously weakest missense subgroup. Concordance remained highest for loss-of-function variants, where predictive tools are most reliable, and lowest for the heterogeneous non-coding ‘Other’ group, consistent with the broader literature on where current annotation tools provide weaker evidence. Discordant classifications were most numerous among single-submitter ClinVar entries (https://www.ncbi.nlm.nih.gov/clinvar/docs/review_status/), in proportion to their share of the benchmark, rather than being disproportionately concentrated there.

Overall, SeverYX offers a transparent (a goal set by the community [21]), extensible framework suitable for both research- and clinical-grade variant interpretation. By unifying disparate predictive signals, mitigating inter-tool inconsistencies, and supporting continuous integration of new computational evidence, it provides a scalable approach to variant severity classification. As variant-effect prediction continues to advance, particularly with deep learning models that integrate multilayer biological information, a framework like SeverYX, built to absorb new calibrated predictors without architectural change, will be well positioned to keep pace, provided its thresholds are periodically revisited as reference datasets and functional assays improve.

## Limitations

Several limitations should be acknowledged. Severity assignments inherit the quality and consistency limitations of upstream annotation tools: splicing predictors show substantial discordance for exonic and synonymous splice-disruptive variants [14], and annotation engines such as ANNOVAR, SnpEff and VEP disagree on HGVS [22] representation and coding-impact classification, particularly for loss-of-function calls [13], discrepancies that propagate into the apparent SeverYX to ClinVar concordance, especially for borderline or multi-transcript variants. Predictive resolution is also weakest for non-coding and regulatory variants, mirrored here by the lower concordance of the ‘Other’ effect category. The ClinVar reference set itself is imperfect: single-submitter entries carry ambiguous or conflicting assertions and accounted for the majority of unexpected classifications, in proportion to their share of the benchmark, though discordance rates were similar across review-status strata, so review status alone does not explain the residual discordance.

A further, more methodological limitation is that because ‘Medium’ is treated as an acceptable outcome for both pathogenic and benign variants, the primary concordance criterion is insensitive to miscalibration within the ‘Medium’ tier. Since the same criterion was also the objective of the threshold search, a configuration that reassigns variants from ‘High’ to ‘Medium’ scores higher on it by construction; the strict concordance figure (90.5%) is reported to bound this effect. Relatedly, several constituent predictors, and the AM/REVEL cut-offs adopted here, were themselves trained or calibrated on ClinVar- or HGMD-derived labels, so reported concordance should be read as an upper bound rather than an independent validation [9,12,23], and because the threshold search was evaluated on the same ClinVar benchmark used to report performance, these estimates are in-sample; prospective evaluation against a later ClinVar release remains necessary.

## Availability of Data and Materials

The ClinVar GRCh38 VCF release of 07 Jan 2024 used to construct the benchmark is available from the NCBI ClinVar archive (https://ftp.ncbi.nlm.nih.gov/pub/clinvar/vcf_GRCh38/archive_2.0/2024/clinvar_20240107.vcf.gz). The preprocessing pipeline (runme.py), the validation and optimization program (severity_validation.py), the severity threshold configuration (thresholds_config.json) and an example optimization configuration specifying the swept threshold ranges (thresholds_config.optimization_example.json) are available at https://github.com/geneyx/geneyx.severity.

## Acknowledgements

This work was carried out as part of Geneyx Genomex’s internal research and development program; it did not receive any specific grant from funding agencies in the public, commercial, or not-for-profit sectors.

## Author Contributions

Conceptualization: Hagar Mor-Shaked (HMS) and Yaron Mazor (YM). Methodology: Evyatar Mor (EM), YM and HMS. Software: EM. Formal analysis: EM, Eldar Dedic (ED), Suzanne Drury (SD), Lina Zelinger (LZ) and HMS. Writing: EM, HMS and LZ. Review and editing: all authors. Supervision: HMS, YM and LZ. All authors read and approved the final manuscript.

## Conflict of Interest

All authors are employees of Geneyx Genomex LTD, which developed SeverYX and the Geneyx Analysis platform described in this work. The authors declare no other competing interests.

Supplementary information (Supplementary Figures 1–5 and Supplementary Table 1) accompanies this preprint.

