## Supplementary figures and tables for "SeverYX: An ensemble framework for robust variant severity classification"

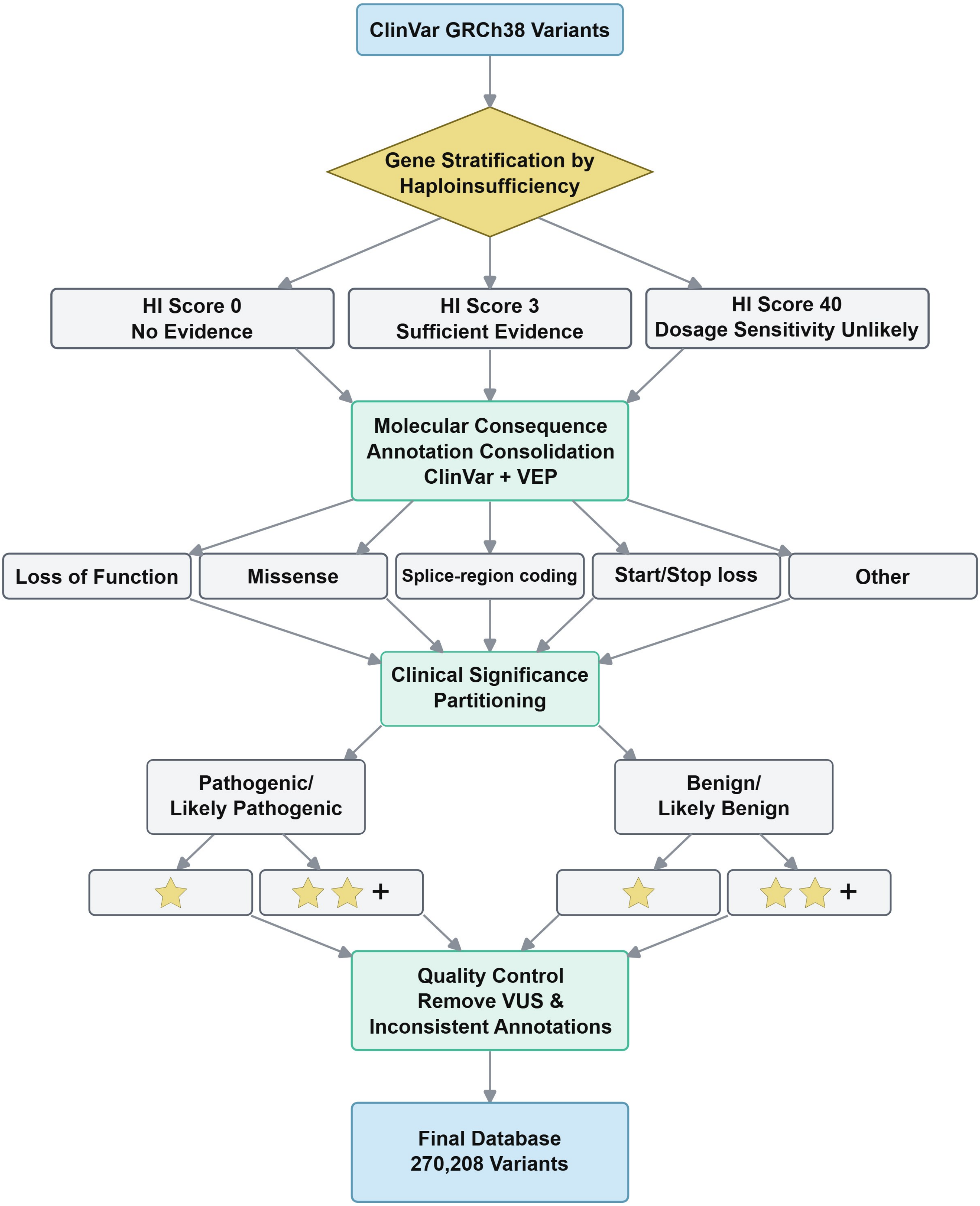


**Supplementary Figure 1. Construction of the ClinVar-based validation dataset.**
Workflow illustrating the generation of the ClinVar benchmark dataset. GRCh38 ClinVar variants were stratified by haploinsufficiency category, consolidated into molecular consequence groups using ClinVar and VEP annotations, partitioned by clinical significance and review status, filtered for VUS and inconsistent annotations, and compiled into the final validation dataset (270,208 variants).

### Supplementary Figure 2

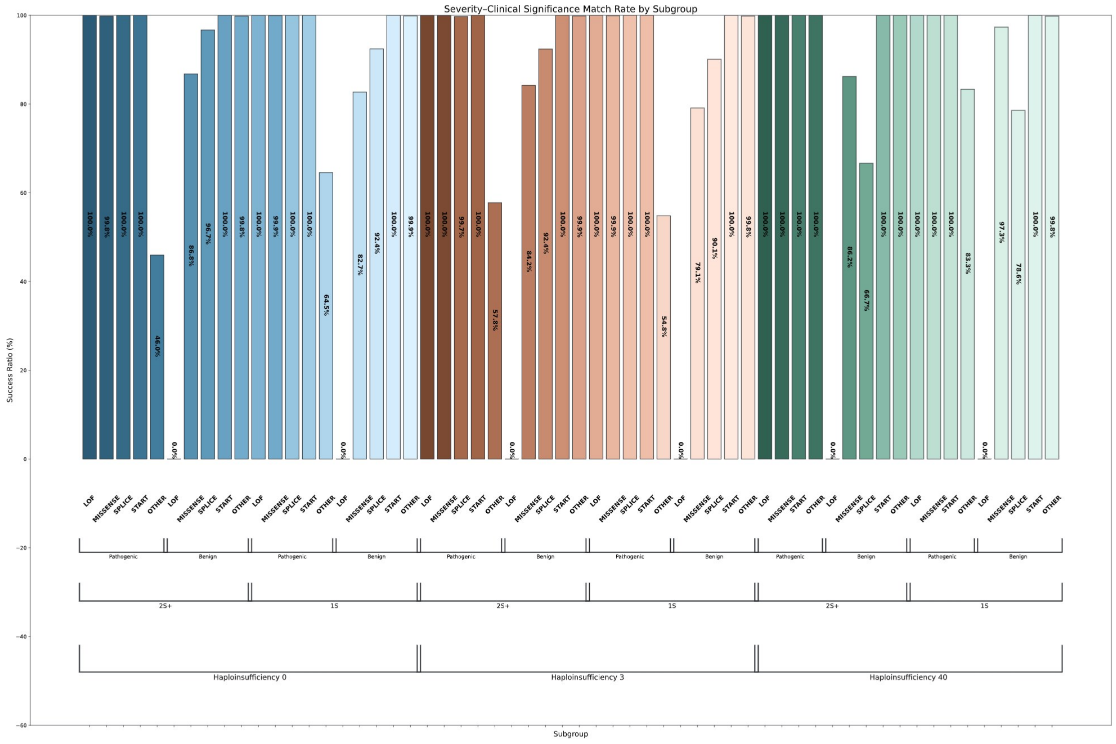


**Supplementary Figure 2.** Expanded subgroup-level concordance between SeverYX and ClinVar.

Expanded view of SeverYX vs. ClinVar concordance for every subgroup defined by haploinsufficiency category (HI-0, HI-3, HI-40), ClinVar review status (one-star vs. two-star-or-better), clinical significance (pathogenic/likely pathogenic vs. benign/likely benign) and variant effect class (loss-of-function, missense, splice-region coding, start/stop, other). Bar heights and the percentage labels printed within each bar give the match rate for that subgroup; a 0.0% bar indicates that no variant in that subgroup received an expected assignment, not that the subgroup is empty.

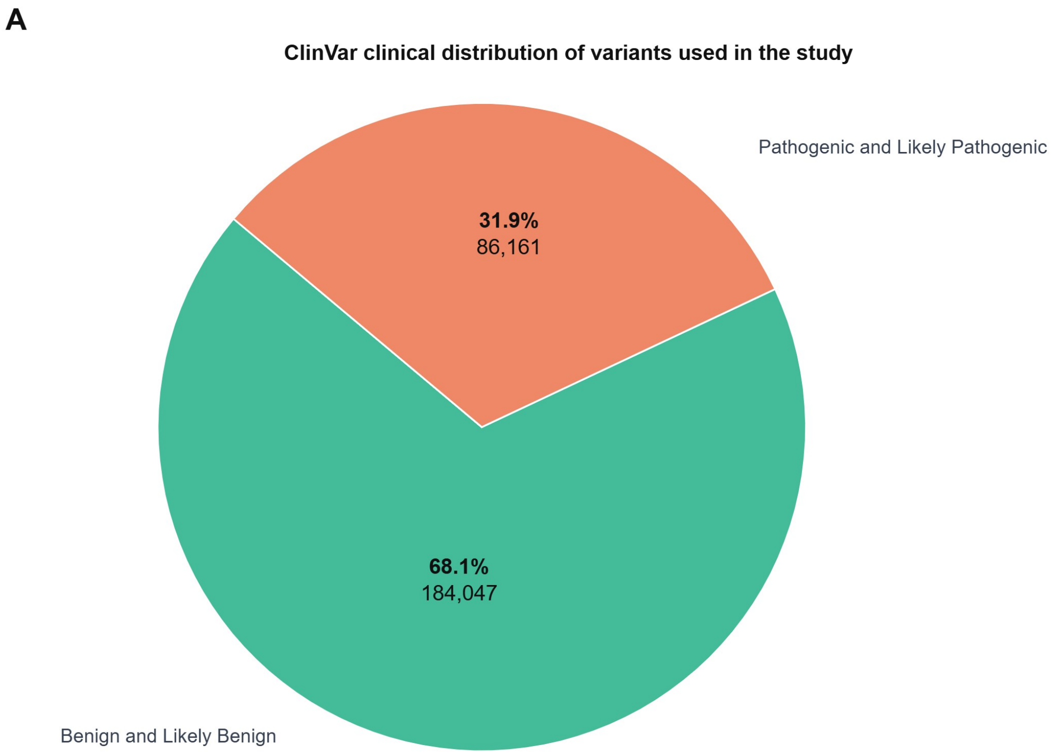


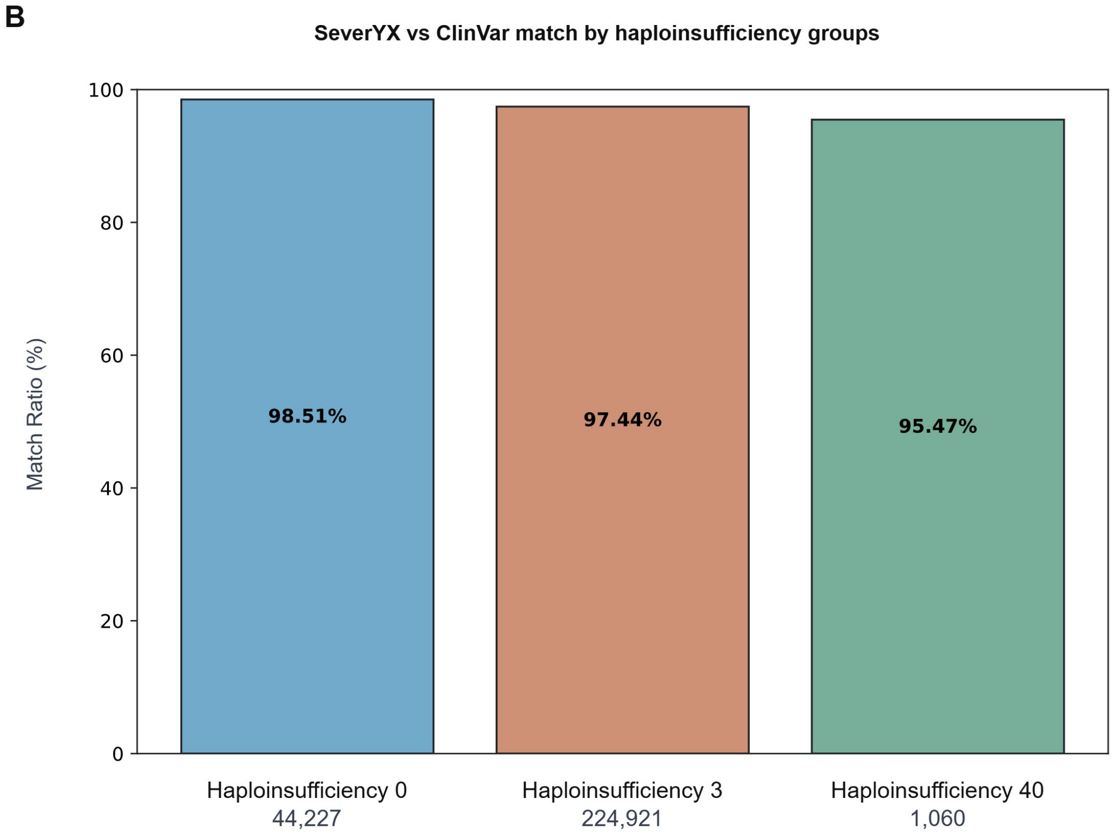


**Supplementary Figure 3.** ClinVar variant composition and SeverYX performance across haploinsufficiency groups.
(A) Clinical significance distribution of all ClinVar variants included in the study. The chart shows the relative proportions of variants annotated as pathogenic or likely pathogenic (86,161; 31.9%) and benign or likely benign (184,047; 68.1%), with corresponding counts displayed for each category. The majority of variants fall within benign or likely benign classifications, reflecting typical ClinVar variant representation. (B) Concordance between SeverYX severity assignments and ClinVar classifications stratified by haploinsufficiency groups. Match ratios are shown for genes with no evidence of haploinsufficiency (HI-0), genes with sufficient evidence of haploinsufficiency (HI-3), and genes for which dosage sensitivity is unlikely (HI-40), with the number of variants in each group listed below the bars. Match rates were 98.51%, 97.44% and 95.47%, respectively; the HI-40 group contains only 1,060 variants and is therefore estimated with wider uncertainty.

### Supplementary Figure 4

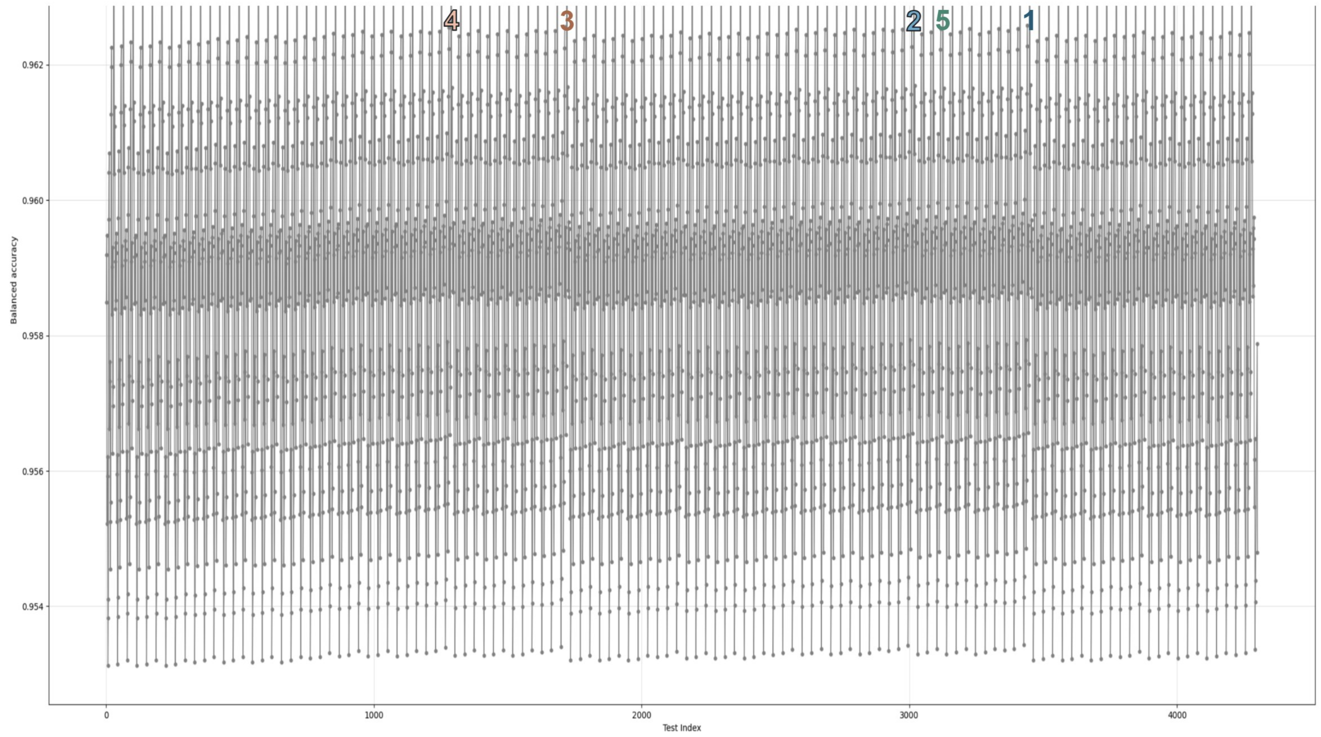


**Supplementary Figure 4.** Balanced accuracy across threshold configurations evaluated during the optimization sweep.
Balanced accuracy of every threshold configuration evaluated during the optimization sweep. Each point is one configuration (4,295 evaluated of 6,912 enumerated), plotted against its configuration index; the five best-performing configurations are highlighted and numbered. No resampling is involved: the dense band reflects the limited total spread of the objective across the search space (balanced accuracy 0.953–0.965). This sweep was run on the rule set that preceded REVEL integration, so these values are not directly comparable with the balanced accuracies reported for the baseline and optimized configurations in the Results.

### Supplementary Figure 5

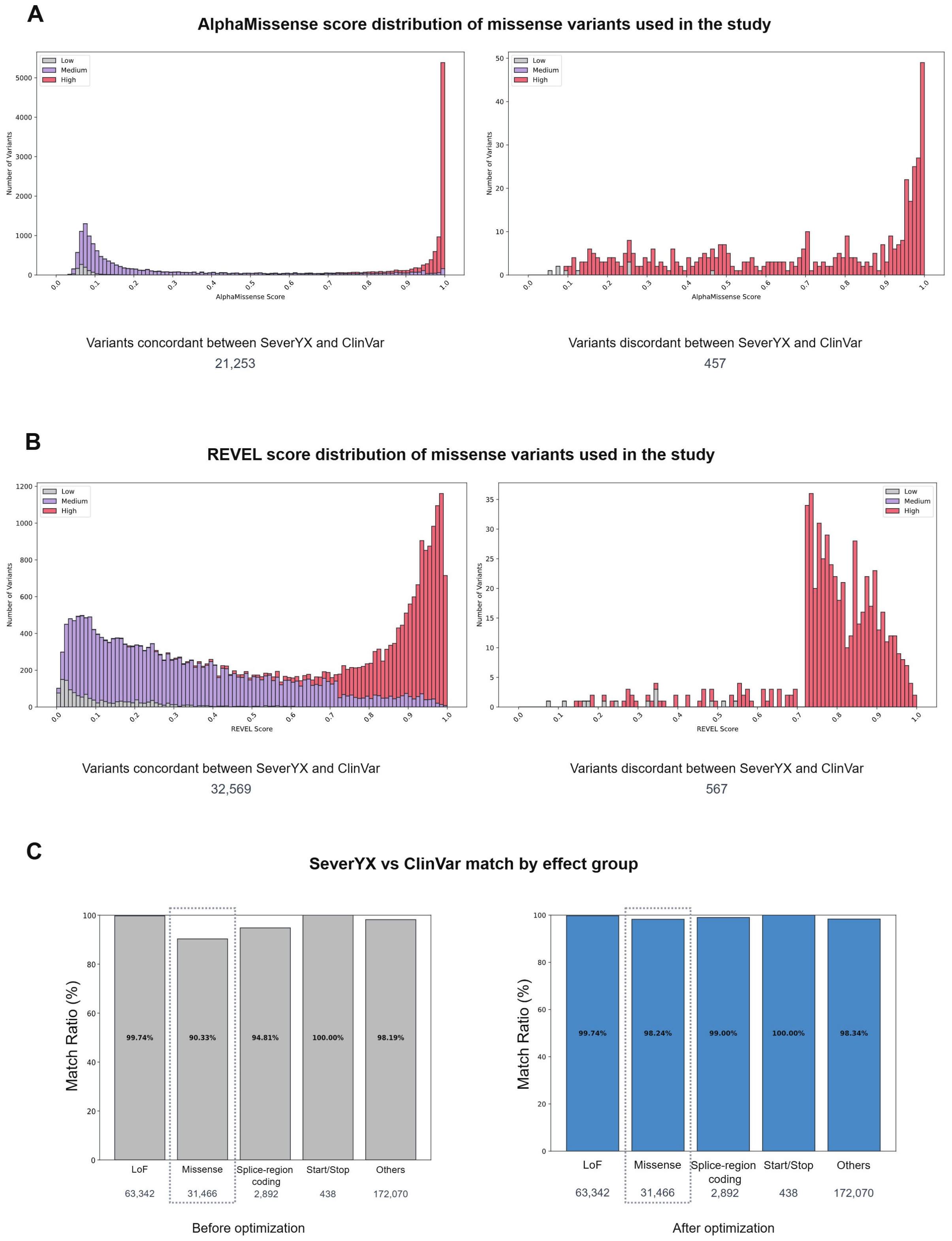


**Supplementary Figure 5.** AlphaMissense and REVEL score distributions for missense variants, and the effect of threshold optimization on match rate by effect group.

(A) AlphaMissense (AM) score distributions for missense variants that were evaluated in this study and where an AM score is available. Left: variants for which SeverYX severity assignments were concordant with ClinVar clinical significance (n = 21,253). Right: variants for which SeverYX and ClinVar were discordant (n = 457).

(B) REVEL score distributions for missense variants that were evaluated in this study and where REVEL score is available, again separated into concordant (left; n = 32,569) and discordant (right; n = 567) sets.

(C) SeverYX vs. ClinVar match rate by variant effect group before (left) and after (right) threshold optimization and the addition of AlphaMissense and REVEL as missense predictors. The missense group (dotted outline) rose from 90.33% to 98.24%, the largest gain of any effect class, while the remaining groups changed only marginally.

**Supplementary Table 1. Combination rule logic**

| | Baseline | | Post optimization | |
|---|---|---|---|---|
| **Variant Type** | **High Requirement** | **Medium Requirement** | **High Requirement** | **Medium Requirement** |
| LoF | 1 of 1 criteria | 1 of 1 criteria | 1 of 1 criteria | 1 of 1 criteria |
| Missense | 4 of 4 criteria | 1 of 8 criteria | 5 of 6 criteria | 1 of 10 criteria |
| Splice-region coding | 1 of 1 criteria | 1 of 1 criteria | 1 of 1 criteria | 1 of 1 criteria |

Criteria are the predictor scores listed in Table 3 for the corresponding variant type.
